# Leaf litter decomposition, microbial respiration and community succession under variable dissolved organic carbon availability in boreal streams

**DOI:** 10.64898/2026.09.25.754356

**Authors:** Matouš Jimel, Lenka Kuglerová, Eric Capo, Judith M. Sarneel, Micael Jonsson

**Affiliations:** Department of Ecology, Environment and Geoscience, Umeå University, Umeå, Sweden; Department of Forest Ecology and Management, Swedish University of Agricultural Sciences, Umeå, Sweden

**Keywords:** dissolved organic carbon, leaf litter decomposition, mesocosm, microbial respiration, microbial communities, boreal streams

## Abstract

Boreal forested streams receive substantial inputs of terrestrial carbon (C) from riparian vegetation in the form of leaf litter, which is processed by litter-associated microorganisms. In addition to these leaf-litter C inputs, boreal streams also receive significant concentrations of dissolved organic carbon (DOC), which may represent an alternative major resource pool for litter-associated microorganisms. As leaf litter quality decreases over the process of decomposition, DOC may increasingly become a more suitable C source for the microbial communities. We investigated how DOC availability, litter type, and microbial succession influenced respiration and decomposition. Alder, birch, and spruce litter were incubated for 65 days in 12 outdoor experimental channels supplied with natural boreal stream water. Higher time-adjusted DOC concentrations were associated with higher respiration and lower relative mass remaining, whereas DOC flux received less support. Respiration and mass loss followed litter-specific trajectories, with birch showing a respiration peak that was not matched by proportional mass loss. Prokaryotic and eukaryotic communities differed among litter types and changed through time but explained process variation only for specific litter types and responses. Our results suggest that DOC can sustain litter-associated microbial activity and partly decouple respiration from litter decomposition, thereby affecting boreal stream C processing.

## Introduction

Terrestrial carbon (C) is a major source of energy for heterotrophic microbial communities in forested headwater streams (Fisher & Likens, 1973; Webster & Benfield, 1986). In the boreal zone, headwater streams are tightly linked to their surrounding catchments and riparian zones, receiving substantial terrestrially derived organic C inputs from the landscape (Laudon et al., 2011; Lidman et al., 2017b). These inputs enter streams both as dissolved organic carbon (DOC) and as coarse particulate organic matter, notably including leaf litter from riparian vegetation (Webster & Benfield, 1986; Laudon et al., 2011; Lidman et al., 2017a, 2017b). Microbial communities, particularly fungi and bacteria, play a central role in processing this organic C (Gessner & Chauvet, 1994; Romaní et al., 2006; Webster et al., 2009). Through decomposition and respiration, microbial processing strongly influences how much terrestrial C is mineralized to carbon dioxide (CO_2_) within streams and how much is transported downstream, which makes streams and rivers important pathways of CO_2_ emission. Because this CO_2_ can derive from both terrestrial inputs and in-stream microbial metabolism, identifying the C sources that support microbial respiration is important for understanding C cycling in boreal headwaters (Cole et al., 2007; Webster et al., 2009; Wallin et al., 2013; Burrows et al., 2017).

Once leaf litter enters streams, it undergoes a sequence of physical, chemical, and biological changes, including leaching of soluble compounds, microbial conditioning, and the subsequent slower breakdown of more structurally complex material (Petersen & Cummins, 1974; Webster & Benfield, 1986; Chauvet, 1987). During these early stages, dissolved compounds released through leaching and microbial processing enter the water column, linking particulate litter inputs to the water-column DOC pool (Bärlocher, 1990; Marks, 2019). The quantity and type of leaf litter entering streams are controlled by riparian vegetation, so variation in riparian plant composition can generate differences in the chemical and structural quality of organic substrates entering boreal streams (Lidman et al., 2017a). Differences among litter types in nutrient content, toughness, and concentrations of recalcitrant compounds influence decomposition dynamics by affecting microbial processing of litter-derived resources (Ostrofsky, 1997; Pastor et al., 2014; Kuglerová et al., 2017). In boreal headwaters, this is particularly relevant because common riparian litter types, including deciduous broadleaf litter and coniferous needles, differ markedly in quality and decomposability (Lidman et al., 2017a). Forestry, land-use change, and climate-driven shifts in riparian vegetation may therefore alter the quality of litter entering streams and its subsequent processing. As a result, the relationship between microbial respiration and litter mass loss is likely to be both litter type-specific and time-dependent, because substrates of differing quality may support microbial activity differently over the course of decomposition (Pastor et al., 2014; Bastias et al., 2022). Litter type is therefore expected to be an important determinant of particulate organic matter processing in boreal headwater streams (Lidman et al., 2017a).

While litter quality can shape the litter-associated microbial community and how this community processes particulate organic matter, these microorganisms are also exposed to DOC in the surrounding water column. Boreal streams receive substantial DOC inputs from organic-rich riparian soils and peatlands, delivered through soil-water and groundwater flow paths, often resulting in high in-stream DOC concentrations (Laudon et al., 2011; Ågren et al., 2008; Bastias & Jonsson, 2024). The availability of this DOC to stream microorganisms may depend not only on its concentration, but also on its hydrological delivery through stream water flow. In many northern catchments, DOC concentrations and exports have increased significantly over recent decades and may continue to change in response to altered climate and land-use regimes (Kritzberg et al., 2020). Much of this DOC is chemically complex, aromatic, and relatively recalcitrant (Ågren et al., 2008; Bastias & Jonsson, 2024). Nevertheless, it represents an important organic C pool available to stream microorganisms, including bacteria in sediments and in the water column (e.g., Berggren et al., 2009).

Recent studies suggest that, for litter-associated microbial communities, DOC in the surrounding water column may provide an additional C source alongside litter-derived C (Pastor et al., 2014; Abril et al., 2019; Bastias et al., 2020, 2022; Bastias & Jonsson, 2024). This use of external C appears to vary among litter types and decomposition stages, with higher DOC use reported for lower-to mid-quality litter and later stages of decomposition (Bastias et al., 2020, 2022). If microbial activity is fueled strictly by litter-derived C, respiration would be expected to remain linked to litter decomposition (i.e., mass loss) over time. Additional DOC could also stimulate microbial processing of litter, producing a priming-like response in which higher respiration is accompanied by greater litter mass loss (Bengtsson et al., 2018; Liu et al., 2026). However, if litter-associated microbes increasingly draw on DOC as litter-derived substrates become less labile, microbial respiration could remain high or increase without a proportional increase in litter mass loss. This would weaken the association between microbial metabolic activity and decomposition of the litter itself. Such decoupling has been observed for birch litter after several months of incubation, when elevated microbial activity was no longer accompanied by greater litter mass loss and the decoupling was more pronounced in streams with higher DOC concentrations (Bastias et al., 2022). However, it remains unclear whether this respiration–mass loss decoupling represents a broader pattern across litter types and whether it varies with DOC concentration, hydrological DOC delivery, or microbial community structure.

Respiration–mass loss relationships may depend not only on litter quality and DOC availability, but also on changes in the microbial communities colonizing the litter. Fungi are often considered the dominant decomposers of leaf litter in streams, particularly because of their ability to colonize and enzymatically degrade plant structural compounds (Gessner & Chauvet, 1994; Duarte et al., 2010). However, bacteria also contribute to litter processing in streams, particularly by using labile organic compounds released during leaching or produced during litter decomposition, and through interactions with fungi during decomposition (Romaní et al., 2006). Both fungal and bacterial communities undergo succession during litter decomposition, meaning that taxa dominating during conditioning and early decomposition stages may differ from those associated with later stages of decomposition (Duarte et al., 2010). Because litter types differ in initial chemistry and structure, these successional trajectories may also vary among substrates as microbial communities respond to changing resource availability during decomposition (Marks et al., 2009; Hayer et al., 2022). These successional changes may alter microbial processing because decomposer groups differ in growth dynamics, substrate use, and extracellular enzyme activity (Romaní et al., 2006). Therefore, litter-specific successional changes in these communities could help explain variation in respiration and mass loss through time and among litter types, although how strongly community structure is linked to these process rates has yet to be determined.

Taken together, evidence from studies of litter quality, DOC use, and microbial succession suggests that litter-associated microbial respiration and litter decomposition may respond differently to DOC availability, litter type, and microbial succession. Distinguishing these responses requires assessing whether DOC availability is associated with respiration and litter mass loss across litter types and decomposition stages, and whether litter-associated microbial community structure helps explain process variation consistent with a shift from decomposition of litter material toward greater use of DOC.

In the present study, we used replicated outdoor experimental stream channels supplied with stream water (Myrstener et al., 2023) to evaluate how DOC availability, including hydrological DOC delivery manipulated through discharge, and litter type were associated with litter mass loss and litter-associated microbial respiration. On five occasions across 65 days, we quantified relative mass remaining, litter-associated microbial respiration, DOC availability, and prokaryotic and eukaryotic community structure. Because DOC availability can reflect both concentration and hydrological delivery, we evaluated DOC concentration and discharge-based DOC flux as alternative representations of DOC availability.

We hypothesized that 1) higher DOC availability, represented by DOC concentration and alternatively by discharge-based DOC flux, would be associated with higher litter-associated microbial respiration. We further expected that DOC availability could either be associated with lower relative mass remaining, consistent with stimulation of litter decomposition, or weaken the relationship between respiration and mass loss if microbes increasingly used DOC as an alternative C source. We further hypothesized that 2) respiration and mass loss would differ among litter types and show different patterns over time, because litter types differ in quality and may therefore differ in when litter-derived substrates become less labile and DOC availability starts to increasingly support microbial respiration. Finally, we hypothesized that 3) prokaryotic and eukaryotic communities would show litter-specific successional trajectories, and that these community differences would explain additional variation in litter-specific respiration and mass loss.

## Materials and methods

### Study site

The study was performed in a fluvial facility within the Krycklan Catchment Study (KCS) area in northern Sweden (64°14′ N, 19°46′ E; Laudon et al., 2021). Krycklan is a well-established boreal research catchment located near the town of Vindeln, 50–60 km northwest of Umeå, and is characterized by forests, mires, streams, and lakes typical of northern boreal landscapes (Laudon et al., 2021). The area has a cold, snow-influenced boreal climate, with a mean annual air temperature of approximately 1.8 °C, mean January and July temperatures of −9.5 °C and 14.7 °C, respectively, mean annual precipitation of 614 mm, and mean annual runoff of 311 mm (Laudon et al., 2013; Teutschbein et al., 2015). In the study, stream C8 (“Fulbäcken”) was used as the water source.

### Study design and DOC availability gradient

The fluvial facility consists of 12 artificial stream channels supplied with continuously pumped stream water via a 3000-L collection tank and four 1000-L distribution boxes, each feeding three channels through outlets (Laudon et al., 2021; Myrstener et al., 2023). The outlets allow moderated water flow (up to 1 L s⁻¹) into 15-m-long channels that are 20 cm wide. Water was routed from the collection tank to the four distribution boxes through PVC connections, ensuring mixing and the same water chemistry entering all channels. Each distribution box fed three channels. Water flow was adjusted to create a gradient in discharge and hydrological DOC delivery by altering the outlet opening and the slope of each channel triplet. Based on discharge measurements during the experiment (see below), flow ranged from 0.04 to 0.84 L s⁻¹ across channels and retrieval dates. Each channel contained a thin layer of sand and semi-randomly placed pebbles, cobbles and small rocks to mimic natural stream substrates. Additionally, each channel had a shading tarp (70% shading) fixed on top to mimic forest canopy shading and to limit growth of filamentous algae (Myrstener et al., 2023).

### Litter bags and litter types

Litter bags were constructed from 6 × 12 cm nylon mesh (250 µm) and heat-sealed into a tetrahedral shape. Each bag contained either 1.0 g of Norway spruce (*Picea abies* (L.) H. Karst) litter, 1.0 g of birch (*Betula* sp.) litter, or 1.3 g of grey alder (*Alnus incana* (L.) Moench) litter, with the greater alder mass used to account for its expected higher mass loss. The three litter types were selected to represent a gradient from relatively recalcitrant Norway spruce to relatively labile grey alder, while also representing common native tree species in the study area. All litter was air-dried at room temperature before bag construction, although spruce needles retained more moisture than alder and birch leaves at the time of weighing.

In total, 360 litter bags were prepared, representing five retrieval dates (2, 9, 22, 50, and 65 days after deployment), three litter types, and 12 channels. Each combination of retrieval date, litter type, and channel was duplicated, with one bag used for respiration measures and the other for mass loss and DNA analyses. Litter bags were deployed in each channel in five retrieval-specific batches, one for each retrieval date, and secured using cable ties attached to a central chain within each channel to hold the litter bags under water. The batches were spaced 50 cm apart along the channel. Within each batch, the two bags of each litter type were placed immediately adjacent to one another, forming one pair per litter type. These litter-type pairs were then arranged approximately 10 cm apart in the order spruce, birch, and alder along the direction of flow to reduce potential effects from species-specific leaching on downstream litter bags, as spruce litter was expected to leach the least, followed by birch litter, and alder litter. Bags were retrieved one batch per retrieval date, beginning with the batch most upstream, closest to the water outlet, and then proceeding downstream.

### Data collection

#### Temperature, light, discharge, water depth and point velocity

In-stream water temperature and light intensity were recorded at 15-min intervals using submerged HOBO Pendant temp/light loggers (Onset, UA-002-64) from 26 August 2024 until the end of the experiment on 30 October 2024. One logger was deployed in each channel and positioned beneath the shading tarp, approximately in the middle of the section where the litter bags were attached, corresponding to the vicinity of the third retrieval batch. Discharge was measured at the outlet of each channel on each retrieval date by timing how long it took to fill a 1.3-L container with channel water and converting this value to discharge (L s⁻¹). A channel-scale DOC flux metric was then calculated as the product of discharge and DOC concentration (discharge × concentration), using measured or interpolated DOC values. On each retrieval date, water depth (cm) and in-channel water velocity (m s⁻¹; Model 801 EM Flow Meter) were measured at five points per channel, spread across the section where the litter bag batches were placed.

#### Mass remaining measurements

Upon retrieval, the bag designated for relative mass remaining and microbial community analysis was kept in a cooling box at approximately 4 °C, frozen, and subsequently freeze-dried using a ScanVac CoolSafe Superior Touch freeze dryer. Birch and alder litter were freeze-dried for a fixed period of 7 days, whereas spruce litter required repeated freeze-drying cycles until mass stabilized. After drying, the litter was removed from the nylon bag and weighed to the nearest 1 mg. Mass remaining was calculated relative to initial post-leaching dry mass (i.e., dry mass after 2 days of field incubation), to exclude mass loss that can occur purely through abiotic processes over the first 24 hours (Petersen & Cummins, 1974). This litter was later used for microbial community analysis.

#### Respiration measurements

At each retrieval, all litter remaining in the bag designated for respiration measurements was placed in a 50 mL Falcon tube, which was then filled with oxygen-saturated source-stream water (C8), leaving little to no headspace. Dissolved oxygen concentration was measured before and after a 3-hour dark incubation using a YSI ProODO optical dissolved oxygen instrument (YSI, Yellow Springs, OH, USA). For each retrieval, three blank tubes containing oxygen-saturated stream water only were included to quantify background respiration in the absence of litter. Respiration was calculated as the change in dissolved oxygen concentration over the 3-hour incubation period, corrected for background respiration in the blanks. Rates are therefore reported as blank-corrected oxygen depletion per liter, per gram litter dry mass, and per hour (mg O₂ L⁻¹ g⁻¹ dry mass h⁻¹). To reduce oxygen measurement imprecision due to temperature changes, the stream water was acclimatized to room temperature before measurements started. After incubation, the litter was freeze-dried and weighed to obtain the dry mass used to standardize respiration rates.

#### DOC measurements

Water samples for DOC analysis were collected from each of the four distribution boxes and their associated channel triplets on 28 August 2024, 17 September 2024, and 30 October 2024. For days without measurements, DOC concentration was estimated by linear interpolation between these dates. Samples were collected in 50 mL Falcon tubes without filtration and stored in a cooling box at approximately 4 °C until delivery to the laboratory. DOC concentrations were determined within 24 hours of sampling at the Biogeochemical Analytical Facility (BAF), Umeå University, by high-temperature combustion of acidified, O₂-bubbled water samples at 870 °C, followed by infrared gas analysis on a Formacs HT-I analyzer (Skalar).

### DNA processing and bioinformatics

#### DNA extraction, PCR amplification, purification, pooling, and sequencing

To characterize litter-associated microbial communities, we used 16S and 18S rRNA gene amplicon sequencing to generate joint prokaryotic and eukaryotic inventories (Yeh et al., 2021). The prokaryotic inventories represent bacterial and archaeal taxa, whereas the eukaryotic inventories include fungi together with other microbial eukaryotes. We therefore interpret these datasets as broad prokaryotic and eukaryotic community inventories rather than as a strict bacteria and fungi comparison.

DNA extraction was performed on all 180 litter samples using the DNeasy PowerSoil Pro kit following the standard protocol. Prior to extraction, 0.2 g of each freeze-dried sample was ground into a uniform fine powder in a Vario MM 500 ball mill using 3 mm balls. DNA extracts from each sample, approximately 25 ng, were then used to PCR amplify the V4 region of the 16S and 18S rRNA genes by targeting a 410 to 595 bp fragment using the primers 515Y (5’-GTGYCAGCMGCCGCGGTAA-3’) and 926R (5’-CCGYCAATTYMTTTRAGTTT-3’) from Yeh et al. (2021). Each PCR was performed in a total volume of 25 µL containing 12.5 µL of GoTaq® Green Master Mix (M712), 9.1 µL of nuclease-free water, and 1.25 µL of each primer at 500 nM. The amplification conditions consisted of an initial denaturation at 95 °C for 2 min, followed by 30 cycles of 45 s at 95 °C, 45 s at 50 °C, and 1 min 30 s at 68 °C, followed by a final 5 min extension step at 68 °C. The molecular inventories were multiplexed using 400 possible molecular barcode combinations with 20 forward and 20 reverse tagged primers. The PCR products were pooled at volumes ranging between 1.5 and 3 µL and purified using the QIAquick PCR Purification kit (Qiagen, Carlsbad, CA, USA). Library preparation using Illumina TruSeq PCR-free chemistry was performed at SciLifeLab Uppsala, and paired-end 2 × 300 bp sequencing using a P1 600 NextSeq 2000 Illumina instrument was performed at SciLifeLab Stockholm.

#### Bioinformatics and ASV assignment

Raw sequence reads were demultiplexed using cutadapt v4.4 (Martin, 2011) with the parameters -e 0.14 --no-indels. Reads were split into prokaryotic- and eukaryotic-assigned reads using the bbsplit function in BBMap (Bushnell, 2014) with the parameters usequality=f qtrim=f minratio=0.30 minid=0.30 pairedonly=f and the reference databases SILVA_132_and_PR2_EUK.cdhit95pc.fasta and SILVA_132_PROK.cdhit95pc.fasta. Prokaryotic-assigned reads were processed as paired reads using the DADA2 pipeline (Callahan et al., 2016). Filtering was performed using filterAndTrim with truncLen=c(230,200), maxN=0, maxEE=c(2,2), truncQ=2, and rm.phix=TRUE, followed by learnErrors, dada, mergePairs, makeSequenceTable, and removeBimeraDenovo. Eukaryotic-assigned reads were processed using forward reads only, with filterAndTrim parameters truncLen=230, maxN=0, maxEE=2, truncQ=2, and rm.phix=TRUE, followed by learnErrors, dada, makeSequenceTable, and removeBimeraDenovo. Taxonomic identification was performed using assignTaxonomy against the SILVA database for prokaryotic ASVs (NR99 v138; Quast et al., 2013) and the PR2 database for eukaryotic ASVs (version 5.0.0 SSU; Guillou et al., 2013). Within the eukaryotic inventory, reads belonging to ASVs assigned to Fungi were classified as fungal-assigned reads.

Before cleaning, 99,950 prokaryotic ASVs were obtained for a total of 19,626,910 reads, and 7,177 eukaryotic ASVs were obtained for a total of 4,556,508 reads. After retaining the 152 biological samples shared between the prokaryotic and eukaryotic inventories, the respective tables contained 97,242 ASVs and 19,012,486 reads and 6,773 ASVs and 4,427,929 reads.

Blank-associated ASVs were removed when their maximum relative abundance in any PCR blank was more than 10% higher than their maximum relative abundance among the biological samples. Blank correction removed 561 prokaryotic ASVs, accounting for 1,129,097 reads, and 27 eukaryotic ASVs, accounting for 7,817 reads. Prokaryotic ASVs classified as non-bacterial assignments (496 ASVs, 6,752 reads), chloroplast-assigned sequences (4,508 ASVs, 1,214,890 reads), mitochondrial-assigned sequences (2,974 ASVs, 172,983 reads), or bacterial ASVs unassigned at the class level (3,337 ASVs, 45,979 reads) were removed. Eukaryotic ASVs classified as animal-associated assignments (722 ASVs, 96,386 reads), plant- or plastid-associated assignments (408 ASVs, 1,706,955 reads), or other non-target or uncertain assignments (1,047 ASVs, 25,608 reads) were removed.

ASVs representing fewer than 0.001% of the reads remaining after blank correction and taxonomic filtering were then removed separately from the prokaryotic and eukaryotic inventories. This corresponded to a minimum total abundance of 164.4 reads in the prokaryotic inventory, removing 72,083 ASVs and 1,595,648 reads, and 25.9 reads in the eukaryotic inventory, removing 3,311 ASVs and 20,992 reads. The final prokaryotic inventory contained 13,283 ASVs and 14,847,137 reads across 152 samples, while the final eukaryotic inventory contained 1,258 ASVs and 2,570,171 reads across the same 152 samples. Rarefaction curves (Fig. S1) were produced for the molecular inventories using the function rarecurve from the vegan R package (Oksanen et al., 2022).

### Statistical analyses

#### Linear mixed-effects models

All statistical analyses were conducted in R version 4.6.0 (R Core Team, 2026). Linear mixed-effects models were fitted using the packages lme4 (Bates et al., 2015) and lmerTest (Kuznetsova et al., 2017). In these models, relative mass remaining and litter-associated respiration were modeled separately. For both responses, models included litter type and incubation duration as fixed effects, with channel identity included as a random intercept to account for repeated observations from the same experimental channel. Incubation duration, measured as days since deployment, was treated as a continuous predictor and centered on its mean before fitting the models to reduce correlation between the linear and quadratic time terms and make the intercept represent the mean incubation duration. A quadratic time term was included in both response models due to expected non-linear change through time. Therefore, the linear time term describes the overall direction of temporal change, while the quadratic term allows the slope to increase, decrease, or level off through time. For respiration, interactions between litter types and both the linear and quadratic time terms were included, allowing each litter type to have a different non-linear respiration trajectory. For relative mass remaining, the model allowed the linear effect of time to differ among litter types, but used a common quadratic time effect for all litter types.

#### DOC availability metrics

Because DOC concentration declined strongly in the supply stream over the course of the experiment, raw interpolated DOC concentration was collinear with incubation duration, which complicates coefficient interpretation (Dormann et al., 2013). To separate the temporal component of DOC variation from deviations around this temporal trend, we first modeled raw interpolated DOC concentration as a function of centered incubation duration and its squared term. The standardized residuals from this model were then used as the DOC predictor in the mixed-effects models. These residuals represent whether DOC concentration was higher or lower than expected for a given point in the experiment, rather than the overall seasonal decline in DOC. We refer to this variable as time-adjusted DOC concentration. This approach is related to residual regression, where the residual component of a predictor is used to represent variation not shared with another collinear predictor (Wurm & Fisicaro, 2014; Iler et al., 2017; González-Medina et al., 2026). However, because DOC concentration was not experimentally manipulated independently of time, this term should be interpreted as a within-time DOC concentration association rather than as a fully independent DOC availability treatment effect. Equivalent models based only on measured, non-interpolated DOC values were also examined. In addition, root mean squared error and mean absolute error analyses were conducted using channels most similar to C8 (low-order headwater streams in the same basin: C4, C5, C6, and C7) to assess interpolation plausibility (Laudon et al., 2007; Ledesma et al., 2018).

Because DOC availability may reflect hydrological delivery as well as concentration, discharge-based DOC flux was evaluated as an *a priori* alternative to DOC concentration. Discharge alone was included as a hydrological comparison to distinguish support for DOC flux from support for flow variation *per se*. Candidate models containing time-adjusted DOC concentration, DOC flux, discharge, and combined DOC–hydrology formulations were compared using AICc under maximum-likelihood fitting. Raw standardized and time-adjusted formulations of discharge and DOC flux were also compared as a sensitivity analysis. Temperature, light intensity, water depth, and point velocity were evaluated separately as additional environmental covariates.

Model selection was conducted by comparing AICc among linear mixed-effects models fitted using maximum likelihood, while also considering parsimony and ecological interpretability (Burnham & Anderson, 2004). Candidate DOC/hydrology and environmental-covariate terms were compared within the selected temporal structure for each response variable. Backward AICc simplification was used as the primary selection procedure. Final models were selected based on AICc, with preference given to lower-AICc models and to simpler models when additional terms did not improve model support. Final parameter estimates were obtained by refitting the selected models using restricted maximum likelihood (REML).

#### Community analyses

Community analyses were conducted separately for the prokaryotic and eukaryotic inventories. Cleaned ASV count data were converted to sample-wise relative abundances, square-root transformed, and used to calculate Bray–Curtis dissimilarities among samples. Community structure was visualized using non-metric multidimensional scaling (NMDS; Kruskal, 1964). Effects of litter type, incubation date, and their interaction were tested using PERMANOVA (adonis2; 999 permutations; Anderson, 2001), and homogeneity of multivariate dispersion by litter type and date was tested using betadisper (Anderson, 2006; Oksanen et al., 2022). Time-adjusted DOC concentration was tested as a predictor of community structure using partial dbRDA after averaging relative abundance profiles across channels within each distribution box, date, and litter type, while controlling for litter type × date and distribution box.

Associations between microbial community structure and the measured process responses (i.e., relative mass remaining and litter-associated respiration) were evaluated using ordination-based and model-based approaches. Response variables were fitted onto three-dimensional NMDS ordinations using envfit. For visualization of NMDS, the response variables were fitted to corresponding two-dimensional NMDS ordinations. In addition, the first five PCoA axes derived from Bray–Curtis dissimilarities were used as microbial community descriptors in litter-specific models of relative mass remaining and respiration, with incubation date, time-adjusted DOC concentration, and channel identity as covariates. Additional explanatory value was quantified as ΔR² relative to models without PCoA axes. The robustness of these results to the dimensional representation of community structure was assessed by repeating the models using three to seven PCoA axes. As a complementary exploratory analysis, partial dbRDA tested unique associations between process variables and community structure after conditioning on litter type × incubation date and channel identity. All community analyses were implemented in R package vegan (Oksanen et al., 2022).

## Results

### Environmental conditions and DOC availability

In-stream temperature declined over time, with the across-channel median of daily median temperatures decreasing from 10.55 °C at the first retrieval date to 3.68 °C at the final retrieval date (Fig. S2a). Channel-level mean temperatures ranged from 7.33 to 7.59 °C across the experiment. Recorded daytime light intensity also declined through autumn, with the across-channel median daytime lux decreasing from 2196 at the first retrieval date to 124 at the final retrieval date (Fig. S2b). Discharge varied among channels and retrieval dates, ranging from 0.04 to 0.84 L s⁻¹ (Fig. S2c). Exploratory inclusion of temperature, light intensity, water depth, and point velocity did not improve model fit for either litter-associated respiration or relative mass remaining, and these variables were therefore not retained in the final models.

DOC availability was characterized using two complementary metrics: measured DOC concentration and calculated channel-scale DOC flux. Measured DOC concentrations in the four distribution boxes ranged from 31.1 to 48.6 mg L⁻¹ over the course of the experiment (Fig. S2d; Table 1; Table S1). DOC concentration declined progressively through time, with mean (± 1 SE) concentrations across boxes decreasing from 48.1 ± 0.2 mg L⁻¹ at the first DOC sampling date to 32.6 ± 0.5 mg L⁻¹ at the final DOC sampling date. Differences among distribution boxes were small relative to the temporal decline, with box-specific mean DOC concentrations ranging from 39.4 to 40.7 mg L⁻¹ across the experiment. Interpolation error, assessed using comparable low-order headwater streams, was low (MAE = 1.80 mg L⁻¹, RMSE = 2.22 mg L⁻¹).

**Table 1.** Environmental conditions at each retrieval date. DOC values from 4 September 2024 and 15 October 2024 are interpolated concentrations. DOC flux refers to channel-scale DOC delivery calculated as discharge × DOC concentration.

| Retrieval date (d/m/y) | Discharge mean (L s<sup>-1</sup>) | Discharge min | Discharge max | DOC mean (mg L<sup>-1</sup>) | DOC min | DOC max | DOC flux mean (mg s<sup>-1</sup>) | DOC flux min | DOC flux max |
| --- | --- | --- | --- | --- | --- | --- | --- | --- | --- |
| 28/08/2024 | 0.32 | 0.08 | 0.64 | 48.12 | 47.90 | 48.60 | 15.52 | 3.79 | 31.07 |
| 04/09/2024 | 0.42 | 0.06 | 0.71 | 45.19 | 44.95 | 45.80 | 18.99 | 2.57 | 32.14 |
| 17/09/2024 | 0.58 | 0.11 | 0.84 | 39.75 | 39.10 | 40.60 | 23.00 | 4.38 | 32.98 |
| 15/10/2024 | 0.29 | 0.04 | 0.71 | 35.08 | 33.89 | 35.59 | 10.09 | 1.42 | 25.25 |
| 30/10/2024 | 0.47 | 0.08 | 0.80 | 32.57 | 31.10 | 33.30 | 15.24 | 2.66 | 26.17 |

Because discharge varied among channels and dates, calculated channel-scale DOC flux also varied, ranging from 1.42 to 32.98 mg s⁻¹ (Fig. S2e). Unlike DOC concentration, DOC flux did not show a consistent temporal decline. Although DOC flux was evaluated as an alternative representation of DOC availability, it received less model support than time-adjusted DOC concentration and was therefore not retained in the final models.

### Leaf litter decomposition

Post-leaching relative mass remaining declined over time for all litter types, although temporal trajectories differed among litter types (Fig. 1a; Table 2). Relative mass remaining differed among litter types (F₂,₁₅₆.₄ = 5.01, p = 0.008) and decreased with incubation duration (F₁,₁₅₆.₇ = 69.98, p < 0.001). The significant positive quadratic time effect indicated that mass loss slowed over time (F₁,₁₅₆.₄ = 4.11, p = 0.044). Temporal trajectories also differed among litter types, as shown by a significant litter type × time interaction (F₂,₁₅₆.₄ = 6.18, p = 0.003). Tukey-adjusted pairwise contrasts indicated that spruce had higher relative mass remaining than alder at the first two retrievals, whereas birch had higher relative mass remaining than alder from day 22 onward and higher relative mass remaining than spruce at the last retrieval dates (50 and 65 days; Table S2).

**Fig. 1.**
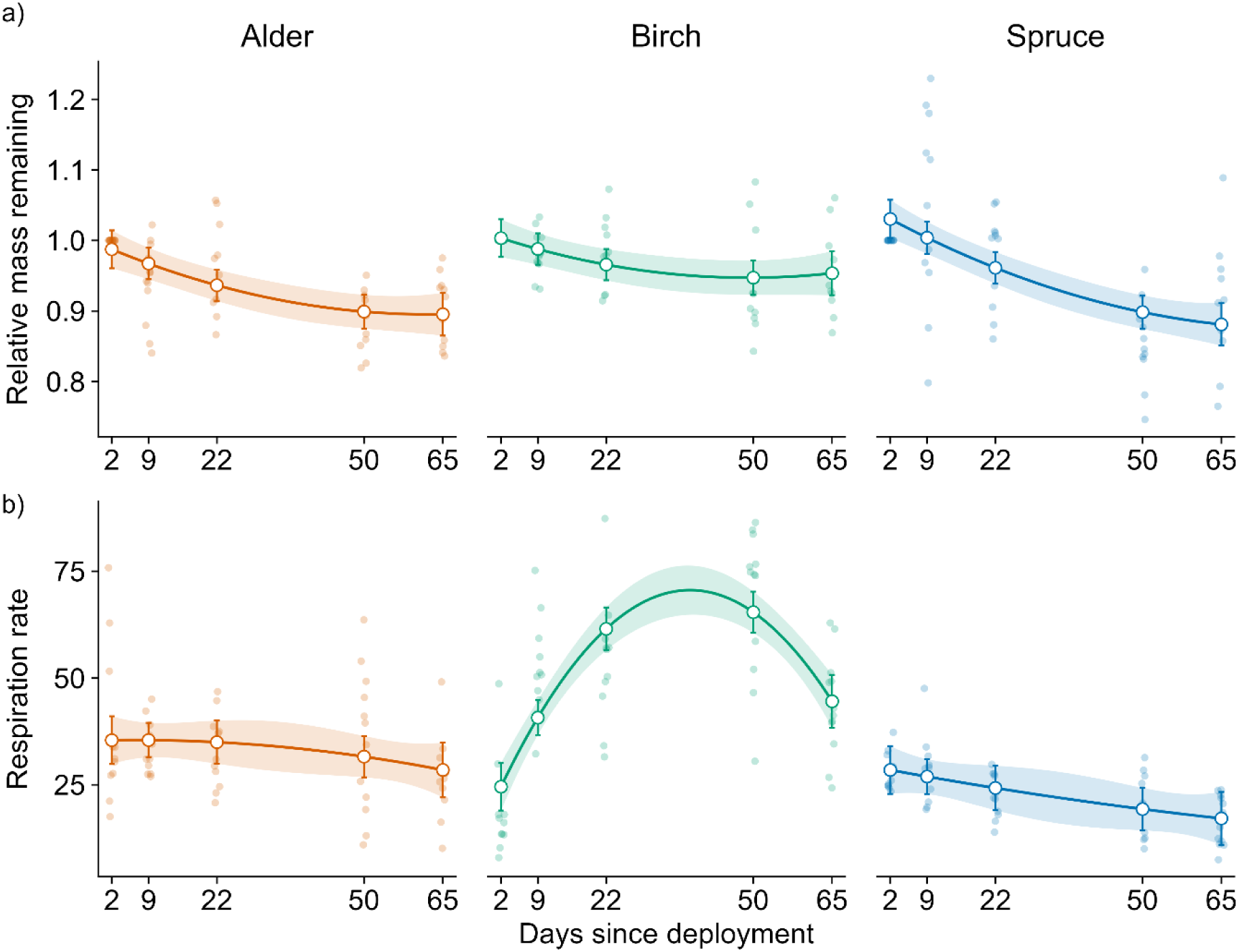
Relative mass remaining (a) and litter-associated respiration rate (b) of alder, birch, and spruce litter through time. Respiration is expressed as blank-corrected oxygen depletion per liter, per gram dry mass remaining, and per hour. Lines and shaded ribbons show fixed-effect model predictions and 95% confidence intervals with time-adjusted DOC concentration held at its mean value, corresponding to DOC concentrations expected from the seasonal DOC trend at each incubation time. Open circles and error bars show predicted values and approximate 95% confidence intervals at the five retrieval dates. Faint points show individual observations

**Table 2.**
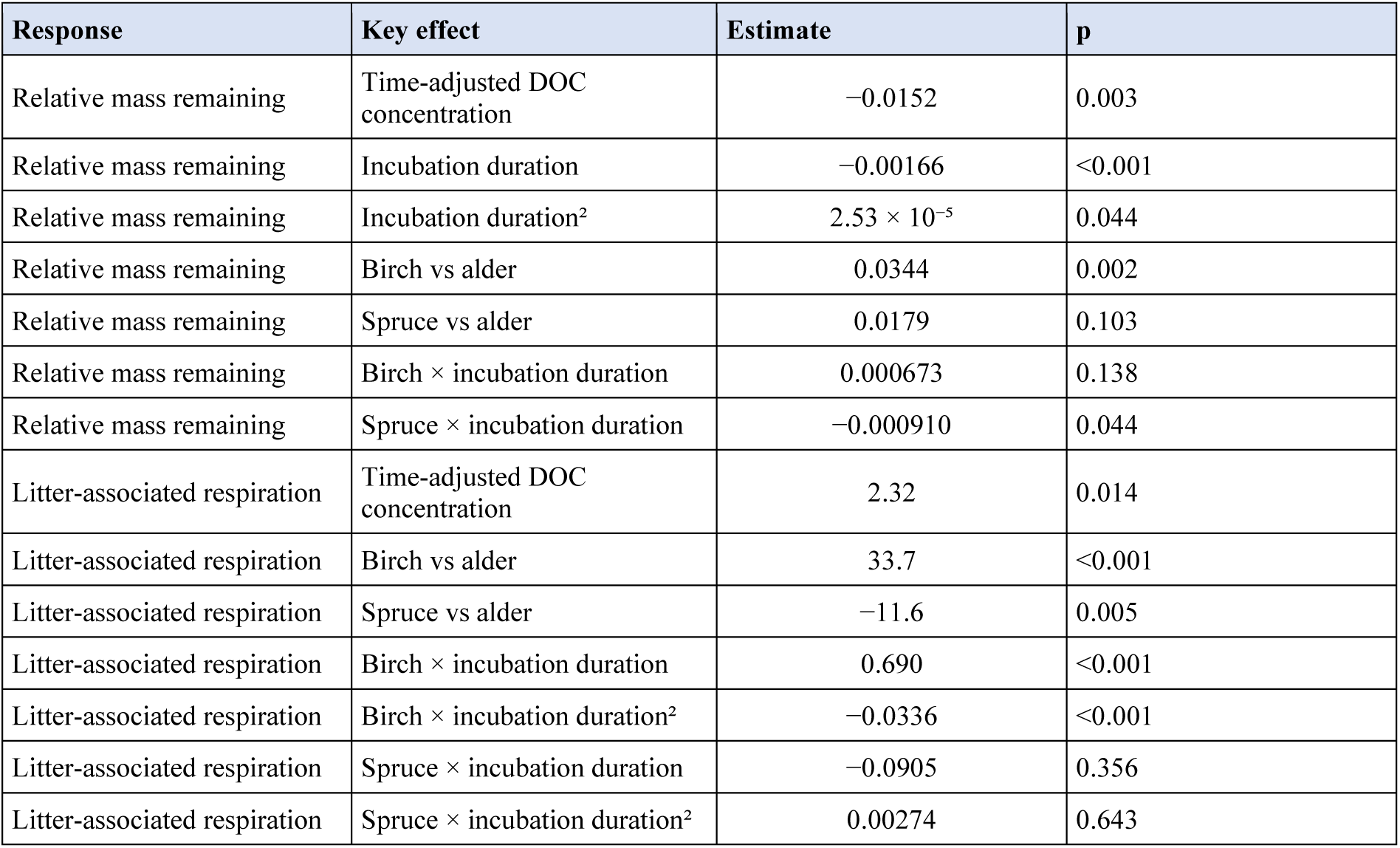
Selected fixed-effect estimates from the final linear mixed-effects models. Respiration was standardized by litter mass remaining. Time-adjusted DOC concentration refers to DOC concentration residualized against incubation duration and incubation duration².

| Response | Key effect | Estimate | p |
| --- | --- | --- | --- |
| Relative mass remaining | Time-adjusted DOC concentration | −0.0152 | 0.003 |
| Relative mass remaining | Incubation duration | −0.00166 | <0.001 |
| Relative mass remaining | Incubation duration<sup>2</sup> | $2.53 \times 10^{-5}$ | 0.044 |
| Relative mass remaining | Birch vs alder | 0.0344 | 0.002 |
| Relative mass remaining | Spruce vs alder | 0.0179 | 0.103 |
| Relative mass remaining | Birch × incubation duration | 0.000673 | 0.138 |
| Relative mass remaining | Spruce × incubation duration | −0.000910 | 0.044 |
| Litter-associated respiration | Time-adjusted DOC concentration | 2.32 | 0.014 |
| Litter-associated respiration | Birch vs alder | 33.7 | <0.001 |
| Litter-associated respiration | Spruce vs alder | −11.6 | 0.005 |
| Litter-associated respiration | Birch × incubation duration | 0.690 | <0.001 |
| Litter-associated respiration | Birch × incubation duration<sup>2</sup> | −0.0336 | <0.001 |
| Litter-associated respiration | Spruce × incubation duration | −0.0905 | 0.356 |
| Litter-associated respiration | Spruce × incubation duration<sup>2</sup> | 0.00274 | 0.643 |

After accounting for litter type and temporal dynamics, time-adjusted DOC concentration explained significant variation in relative mass remaining (F₁,₁₄₄.₄ = 9.25, p = 0.003). Higher time-adjusted DOC concentration was associated with lower relative mass remaining (β = −0.015 ± 0.005 SE, t₁₄₄.₄ = −3.04, p = 0.003).

### Litter-associated respiration

Litter-associated respiration differed significantly among litter types and showed litter-specific temporal trajectories (Fig. 1b; Table 2). Respiration differed strongly among litter types (F₂,₁₅₆.₅ = 68.84, p < 0.001) and varied through time, with significant linear (F₁,₁₅₆.₄ = 6.97, p = 0.009) and quadratic time effects (F₁,₁₅₆.₅ = 26.41, p < 0.001). These temporal patterns differed among litter types, as indicated by significant litter type × time (F₂,₁₅₆.₄ = 38.73, p < 0.001) and litter type × time² (F₂,₁₅₆.₅ = 24.01, p < 0.001) interactions. At the mean incubation duration, birch litter respiration was substantially higher than alder litter (β = 33.72 ± 3.96 SE, t₁₅₆.₃ = 8.51, p < 0.001), whereas spruce litter respiration was lower than alder litter (β = −11.58 ± 4.04 SE, t₁₅₆.₆ = −2.86, p = 0.005). Birch litter respiration showed a pronounced hump-shaped temporal trajectory, with a positive linear interaction (β = 0.690 ± 0.097 SE, t₁₅₆.₃ = 7.12, p < 0.001) and a negative quadratic interaction (β = −0.0336 ± 0.0058 SE, t₁₅₆.₄ = −5.76, p < 0.001). In contrast, spruce litter respiration did not differ significantly from alder litter in either its linear or quadratic temporal trajectory.

After accounting for litter type and temporal dynamics, time-adjusted DOC concentration explained significant variation in litter-associated respiration (F₁,₁₂₂.₀ = 6.29, p = 0.014). Higher time-adjusted DOC concentration was associated with higher litter-associated respiration (β = 2.32 ± 0.93 SE, t₁₂₂.₀ = 2.51, p = 0.014).

### Microbial community structure and taxonomic composition

NMDS analysis and subsequent PERMANOVA showed that litter-associated microbial community structure changed significantly with both litter type and incubation date in both the prokaryotic and eukaryotic inventories (Fig. 2; Table 3). For the prokaryotic community, litter type explained 27.4% of the variation in community structure (R² = 0.274, F = 32.91, p = 0.001), while incubation date explained 11.6% (R² = 0.116, F = 6.94, p = 0.001). For the eukaryotic community, litter type explained 25.1% of the variation in community structure (R² = 0.251, F = 27.21, p = 0.001), while incubation date explained 7.7% (R² = 0.077, F = 4.16, p = 0.001).

**Fig. 2.**
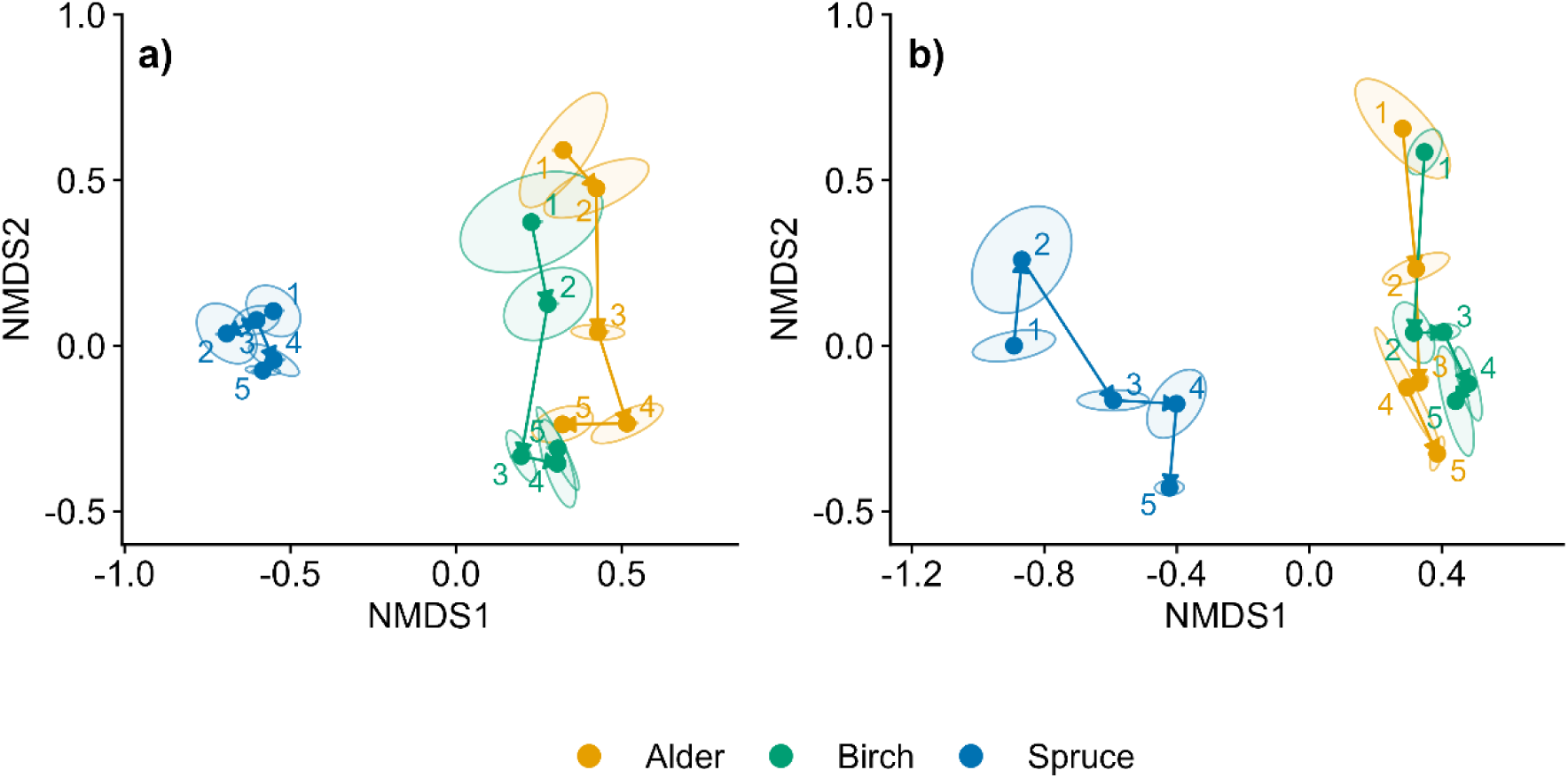
NMDS ordination of eukaryotic (a) and prokaryotic (b) community structure based on Bray-Curtis dissimilarities. Points represent litter-specific centroids for each retrieval date, and arrows connect centroids in chronological order. Ellipses indicate standard errors around centroid positions. Numeric labels denote retrieval order: 1 = 28 August, 2 = 4 September, 3 = 17 September, 4 = 15 October, and 5 = 30 October 2024. Colors denote litter type: alder (yellow), birch (green), and spruce (blue). NMDS stress values were 0.188 for the eukaryotic inventory and 0.157 for the prokaryotic inventory

**Table 3.**
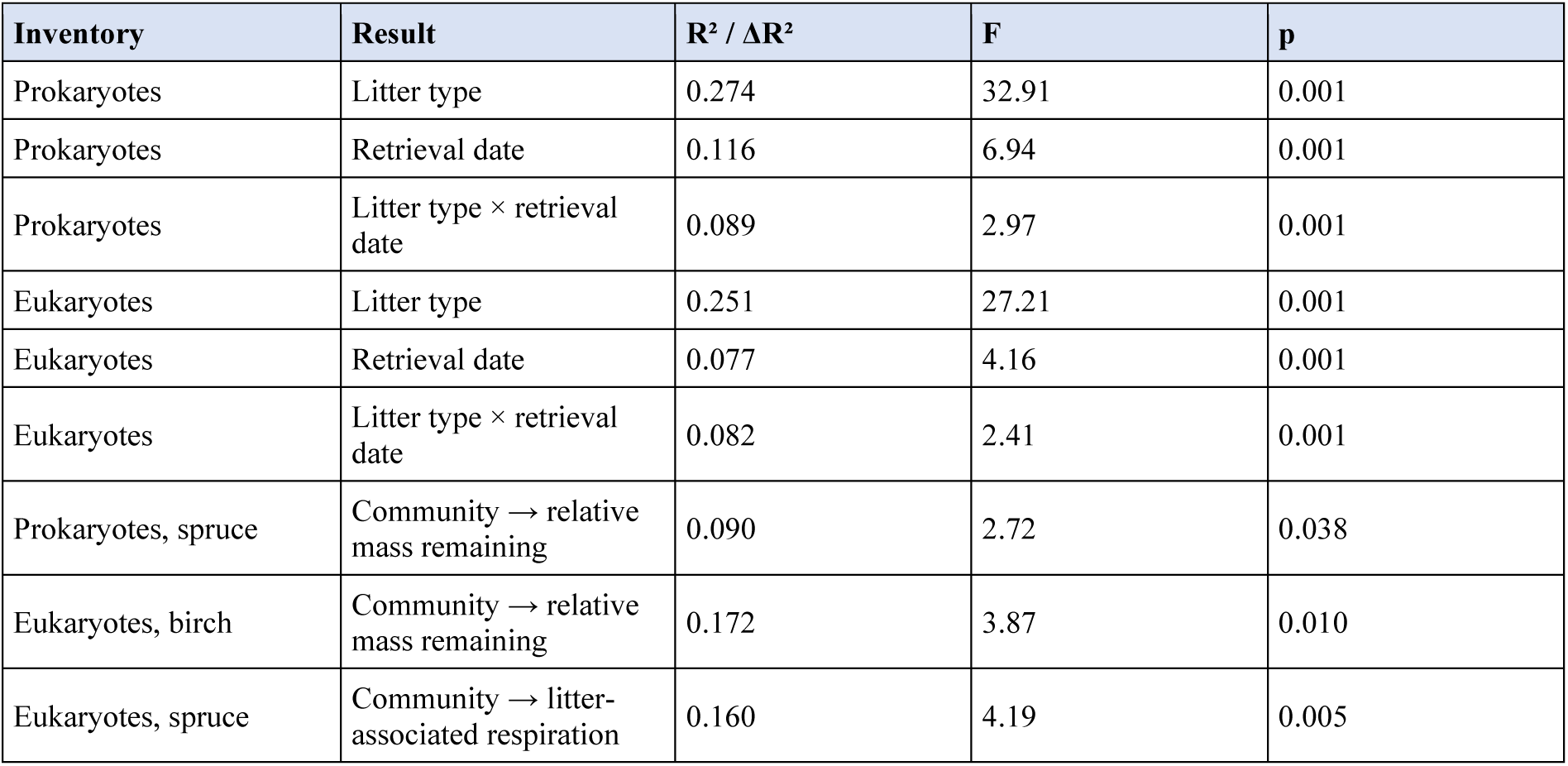
Summary of community structure analyses for prokaryotic and eukaryotic inventories. PERMANOVA rows show effects of litter type, retrieval date, and their interaction on community structure. Community-process rows summarize litter-specific PCoA models testing whether community structure explained additional variation in relative mass remaining or litter-associated respiration after accounting for experimental structure.

| Inventory | Result | R<sup>2</sup> / ΔR<sup>2</sup> | F | p |
| --- | --- | --- | --- | --- |
| Prokaryotes | Litter type | 0.274 | 32.91 | 0.001 |
| Prokaryotes | Retrieval date | 0.116 | 6.94 | 0.001 |
| Prokaryotes | Litter type × retrieval date | 0.089 | 2.97 | 0.001 |
| Eukaryotes | Litter type | 0.251 | 27.21 | 0.001 |
| Eukaryotes | Retrieval date | 0.077 | 4.16 | 0.001 |
| Eukaryotes | Litter type × retrieval date | 0.082 | 2.41 | 0.001 |
| Prokaryotes, spruce | Community → relative mass remaining | 0.090 | 2.72 | 0.038 |
| Eukaryotes, birch | Community → relative mass remaining | 0.172 | 3.87 | 0.010 |
| Eukaryotes, spruce | Community → litter-associated respiration | 0.160 | 4.19 | 0.005 |

Community trajectories through time differed among litter types (Fig. 2; Table 3). PERMANOVA detected significant litter type × incubation date interactions for both the prokaryotic (R² = 0.089, F = 2.97, p = 0.001) and eukaryotic inventories (R² = 0.082, F = 2.41, p = 0.001). Multivariate dispersion also differed among litter types and dates in both inventories, indicating that some group differences reflected variation in within-group heterogeneity, although these effects were generally weaker than the corresponding PERMANOVA centroid differences. Dispersion differed by litter type and date in the prokaryotic inventory (litter type: F = 3.30, p = 0.039; date: F = 3.09, p = 0.013) and the eukaryotic inventory (litter type: F = 9.23, p = 0.001; date: F = 3.39, p = 0.017). Time-adjusted DOC concentration was not associated with prokaryotic (R² = 0.007, F = 0.96, p = 0.400) or eukaryotic community structure (R² = 0.009, F = 0.97, p = 0.419).

Unconditioned envfit vectors showed the broad alignment of respiration and relative mass remaining with NMDS community gradients (Fig. 3). Taxonomic summaries provided a descriptive overview of the dominant groups in the prokaryotic inventory and in the full eukaryotic inventory, including its fungal-assigned subset, across litter types and retrieval dates (Figs. 4–6). The prokaryotic community was dominated by Gammaproteobacteria, Alphaproteobacteria, and Bacteroidia, which accounted for 47.8%, 21.9%, and 17.1% of prokaryotic reads, respectively (Fig. 4). The clearest temporal differences in relative read proportions were observed for Gammaproteobacteria and Bacteroidia, with Gammaproteobacteria contributing proportionally less from the first to the final retrieval (49.6% to 39.2%) and Bacteroidia proportionally more at the final retrieval than at the first retrieval (15.6% to 20.7%). The eukaryotic community was dominated by fungal-assigned reads, which accounted for 83.9% of eukaryotic reads overall, although this pattern differed among litter types (Fig. 5). Fungal-assigned reads represented most eukaryotic reads on spruce and birch litter, accounting for 94.9% and 88.9% of eukaryotic reads, respectively, whereas alder litter had a larger non-fungal eukaryotic component, mainly Ciliophora, Cercozoa, and Gyrista. The fungal-assigned subset was dominated by unclassified Pezizomycotina, Agaricomycetes, Leotiomycetes, and Tremellomycetes, which accounted for 24.4%, 20.5%, 15.0%, and 11.3% of fungal reads, respectively (Fig. 6).

**Fig. 3.**
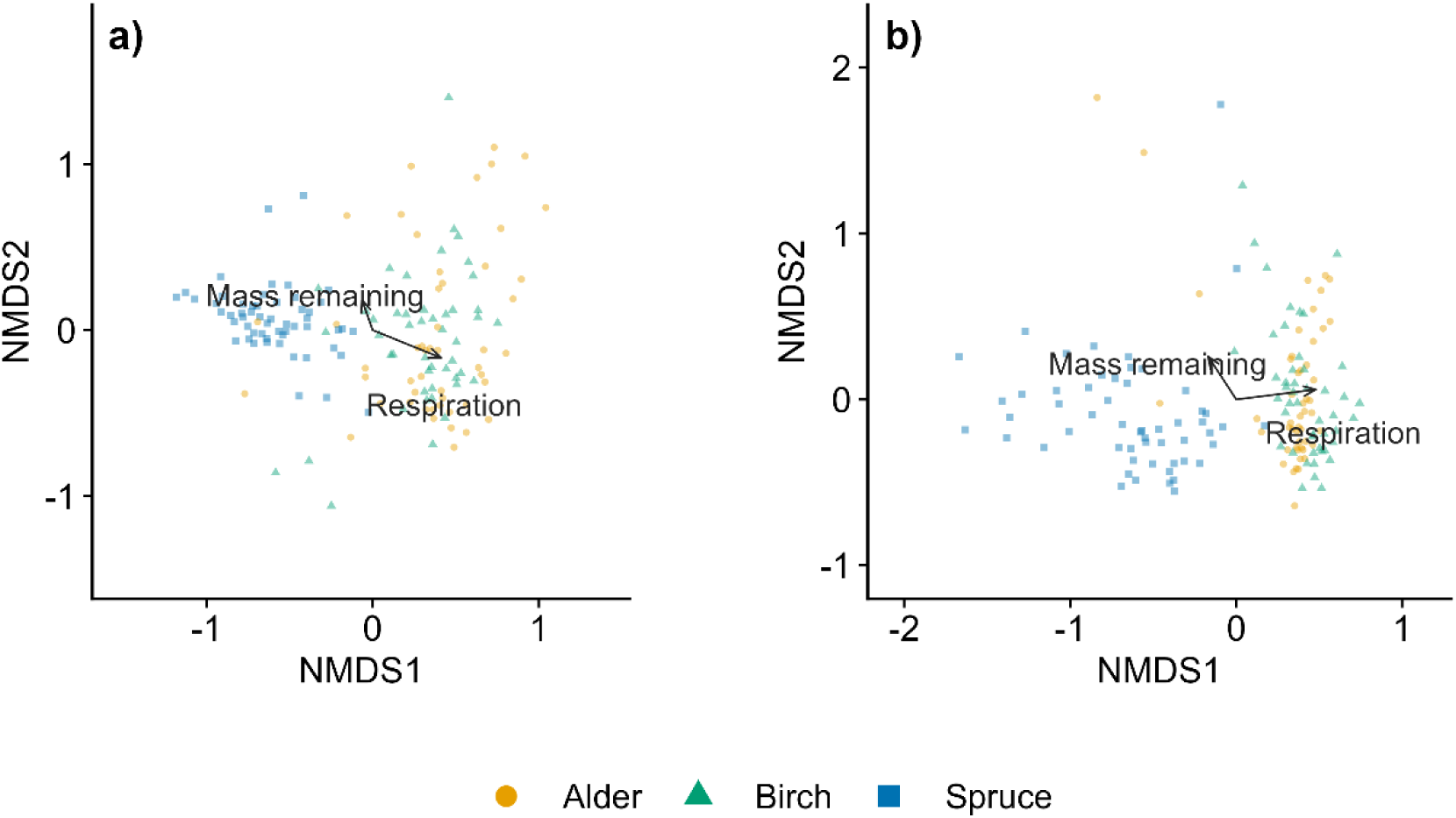
NMDS ordination of eukaryotic (a) and prokaryotic (b) community structure based on Bray-Curtis dissimilarities. Points represent individual litter samples, and colors and shapes denote litter type. Arrows show unconditioned envfit vectors for litter-associated respiration and relative mass remaining, indicating broad alignment with community gradients. These vectors are shown as exploratory visual summaries and do not account for litter type, retrieval date, or channel identity. NMDS stress values were 0.188 for the eukaryotic inventory and 0.157 for the prokaryotic inventory

**Fig. 4.**
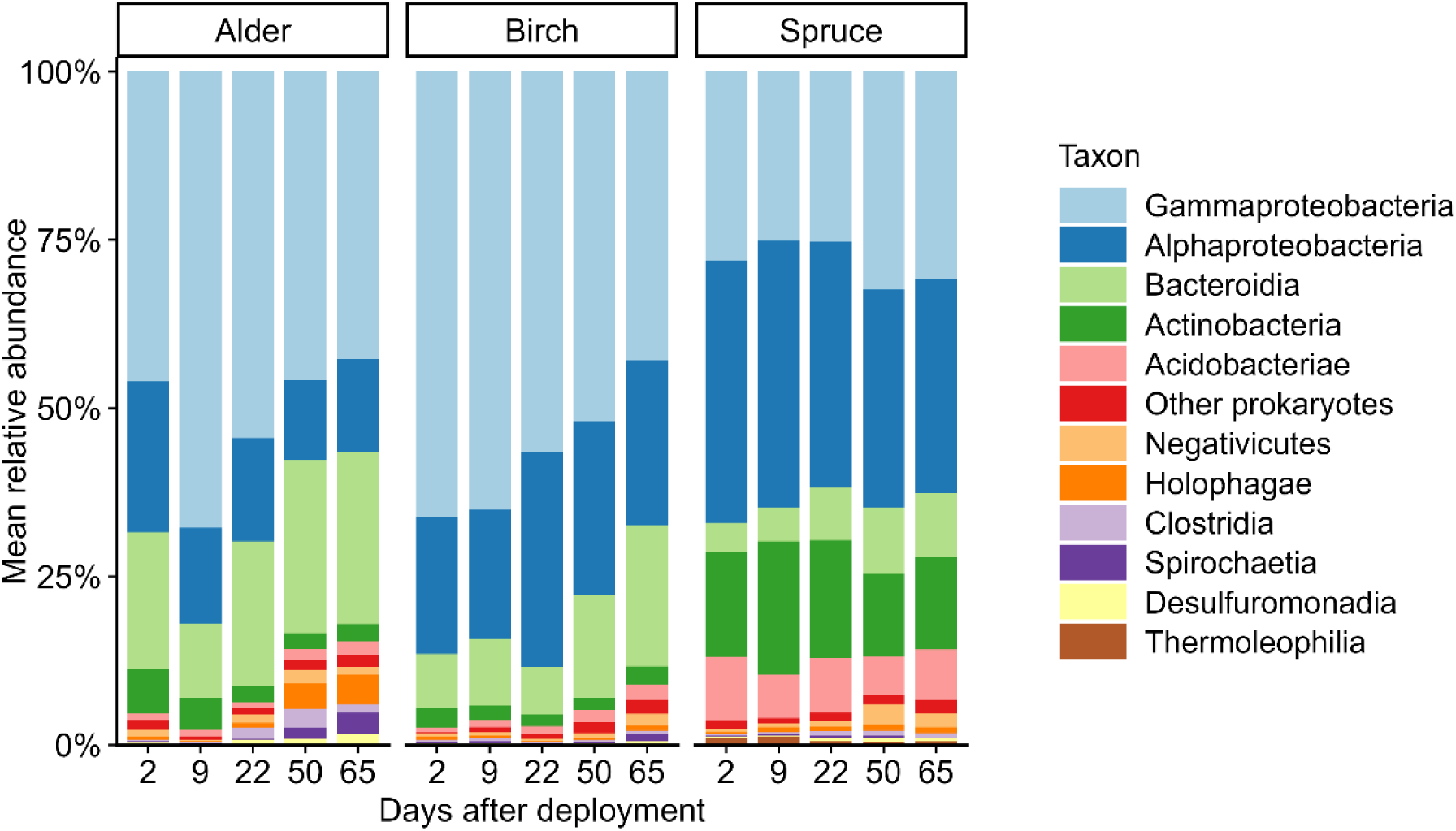
Prokaryotic community composition across litter types and retrieval dates. Bars show sample-level mean relative abundance of dominant prokaryotic classes within each litter type and retrieval date combination. Retrieval dates correspond to 2, 9, 22, 50, and 65 days after deployment

**Fig. 5.**
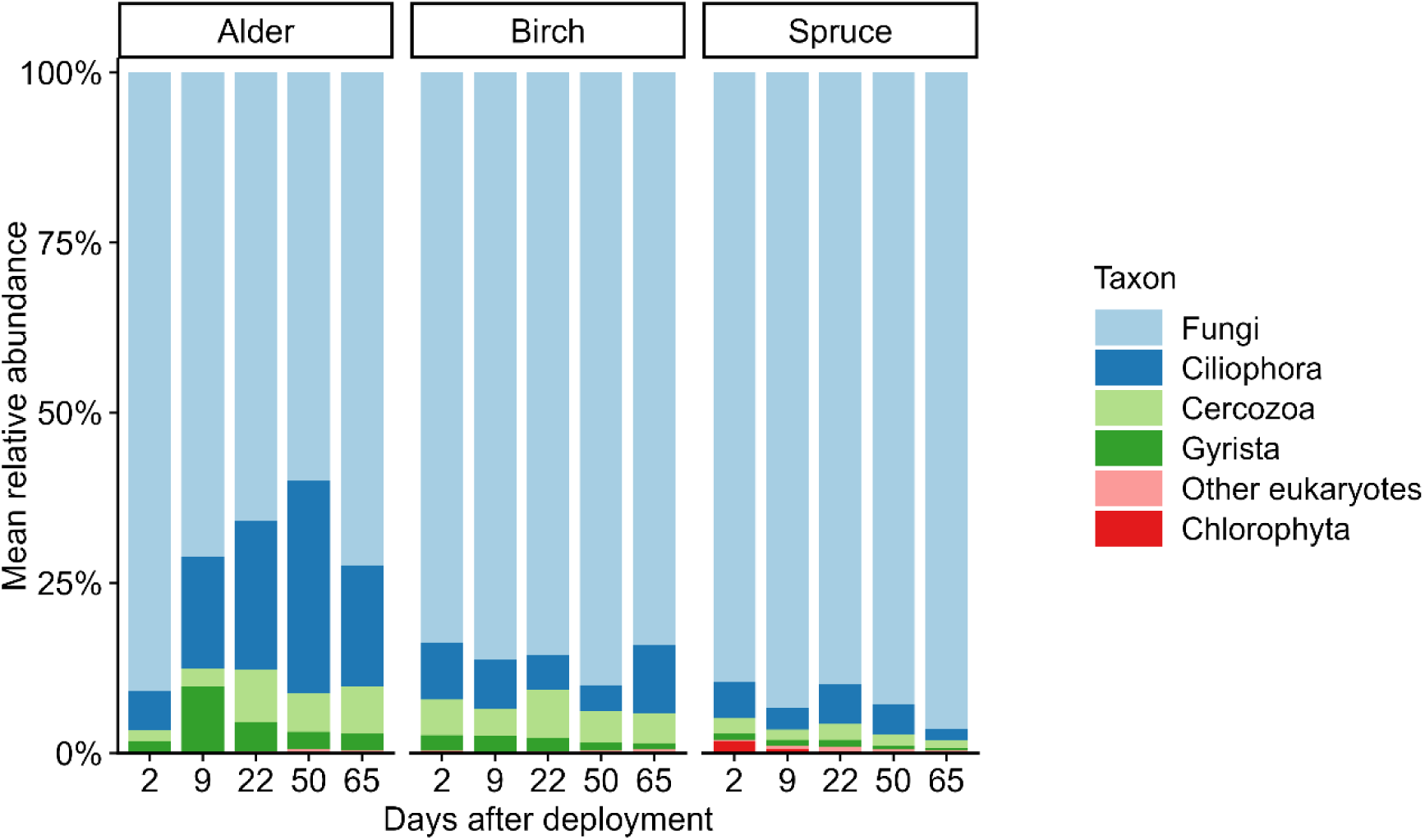
Eukaryotic community composition across litter types and retrieval dates. Bars show sample-level mean relative abundance of broad eukaryotic groups within each litter type and retrieval date combination. Retrieval dates correspond to 2, 9, 22, 50, and 65 days after deployment

**Fig. 6.**
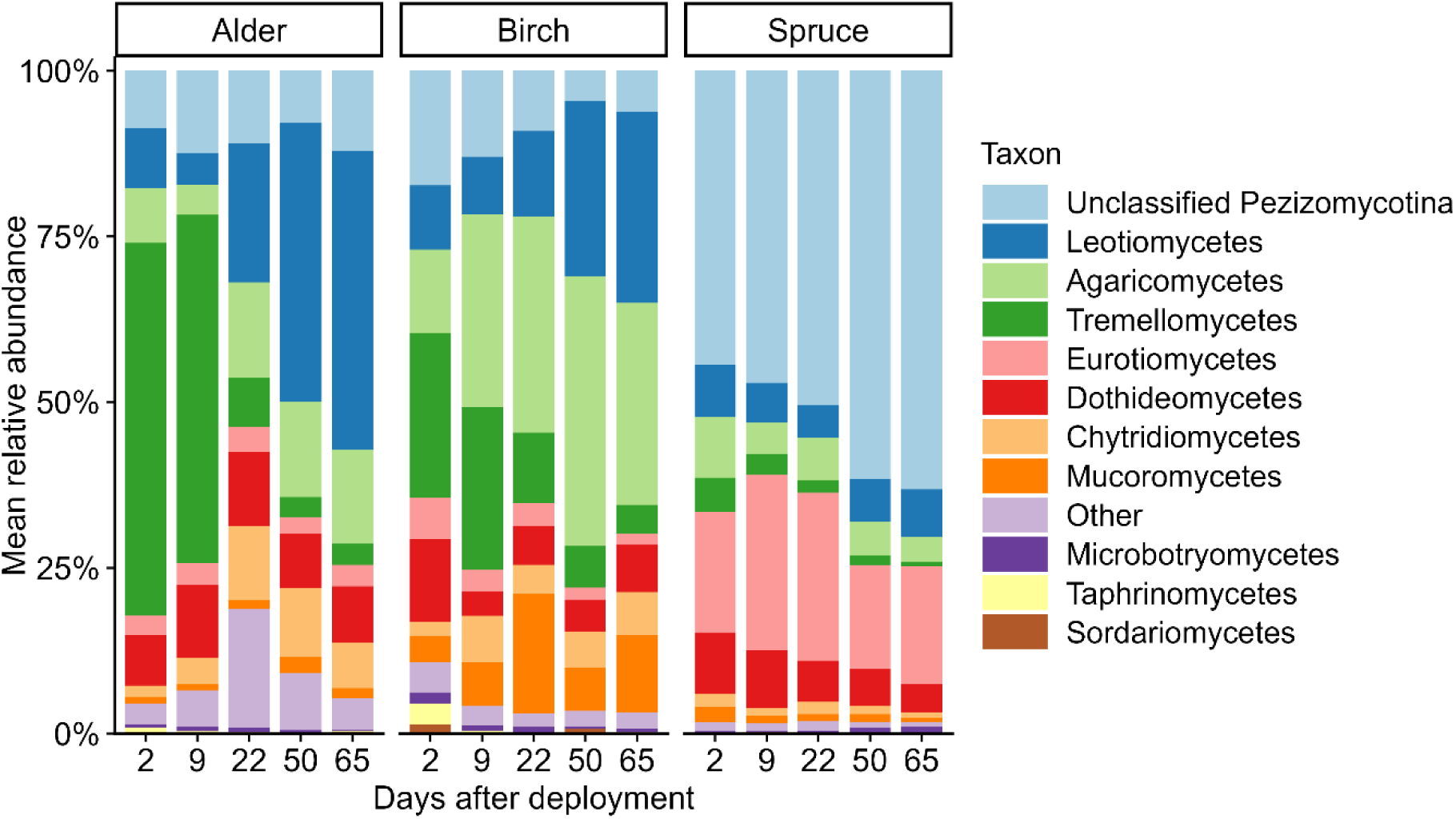
Fungal-assigned subset of the eukaryotic inventory across litter types and retrieval dates. Bars show sample-level mean relative abundance of dominant fungal groups within fungal-assigned reads for each litter type and retrieval date combination. Retrieval dates correspond to 2, 9, 22, 50, and 65 days after deployment. Unclassified Pezizomycotina represents reads assigned to Pezizomycotina but not resolved further to class level

After accounting for incubation date, channel identity, and time-adjusted DOC concentration, litter-specific PCoA models indicated that associations between community structure and process responses were both inventory- and litter-specific. Prokaryotic community structure explained additional variation in relative mass remaining for spruce litter (ΔR² = 0.090, F = 2.72, p = 0.038). Eukaryotic community structure explained additional variation in relative mass remaining for birch litter (ΔR² = 0.172, F = 3.87, p = 0.010) and respiration for spruce litter (ΔR² = 0.160, F = 4.19, p = 0.005).

## Discussion

In this study, litter type, incubation duration, and time-adjusted DOC concentration were the most important factors affecting litter-associated respiration and relative mass remaining in running water. Higher time-adjusted DOC concentration was associated with higher respiration and lower relative mass remaining, suggesting greater microbial activity and litter decomposition, where DOC concentration was higher than expected for a given incubation duration. An important distinction is that respiration responded more strongly than litter decomposition to DOC concentration, creating a discrepancy between the two processes and implying that the increased metabolism may have been supplemented by DOC. Although microbial community structure differed among litter types and changed across time, links between community structure and process rates were detected only for specific litter types and responses. Together, these results suggest that litter-associated microbes may be able to maintain increased metabolic activity through the use of alternative C sources such as DOC.

The weak link between community structure and process rates was notable because litter type and incubation date strongly structured both prokaryotic and eukaryotic communities, whereas links between community structure and respiration or litter decomposition were detected only for specific litter types and responses. This is supported by broader evidence that litter decomposition is strongly regulated by litter traits and resource quality, while microbial community structure may be only weakly linked to measured functions (Gessner & Chauvet, 1994; Gessner et al., 2010; Purahong et al., 2014; Zhang et al., 2019). One explanation for this weak link is functional redundancy, where different assemblages retain similar litter decomposition functions despite taxonomic turnover (Gessner et al., 2010; Schroeter et al., 2022). Alternatively, process rates may have been driven more directly by microbial biomass, physiological activity, enzyme expression, or substrate availability than by community structure itself (Artigas et al., 2009; Purahong et al., 2014), which may help explain why microbial succession in this study was more evident as community turnover, rather than as a consistent predictor of respiration or decomposition.

The taxonomic shifts observed in this study provide one example of such community turnover and are consistent with substrate-driven microbial succession, where changes in the availability of litter-derived compounds during decomposition may favor different microbial groups over time. In the prokaryotic inventory, the higher relative proportion of Bacteroidia and lower relative proportion of Gammaproteobacteria at later dates suggest that the bacterial component of the litter colonizing community changed as litter decomposition progressed, rather than remaining a stable assemblage passively attached to the litter. Bacteroidia belong to a broader group that includes many taxa capable of degrading complex carbohydrates and polymeric organic matter, which is consistent with a role in later processing of plant-derived or microbially modified substrates (Thomas et al., 2011; Fernández-Gómez et al., 2013). Thus, the prokaryotic succession may reflect a shift from early use of readily soluble litter-derived compounds toward later processing of microbial and litter-derived substrates, including substrates generated through fungal activity or litter conditioning (Tláskal et al., 2016). However, because the molecular inventories describe relative taxonomic composition rather than microbial biomass, enzyme expression, or C use, these patterns should be interpreted as consistent with microbial succession rather than direct evidence of the mechanisms controlling respiration or decomposition.

Because microbial community structure was only selectively associated with process rates, the clear mismatch between respiration and mass loss in birch is unlikely to be explained by taxonomic succession alone. This pattern is consistent with previous work on microbial communities associated with birch litter in boreal streams (Bastias et al., 2022) and suggests that birch may have initially supported high microbial activity through relatively accessible litter-derived compounds. After these labile compounds declined, DOC in the surrounding water could have helped maintain and even increase respiration without a corresponding increase in litter decomposition. This interpretation is consistent with studies showing that litter-associated microbes can use stream water C and nutrients during litter decomposition, and that this use differs among litter types and decomposition stages (Gessner & Chauvet, 1997; Bastias et al., 2020; Bastias & Jonsson, 2024).

The interpretation that litter-associated microbes use DOC has an important methodological as well as mechanistic implication, because respiration measured from microbes associated with litter is not necessarily respiration of litter-derived C (Gessner & Chauvet, 1997). Decomposing litter functions both as a substrate and as a microbial habitat. Mass loss measures the reduction in litter material, whereas respiration measures microbial metabolism occurring on or around that material, and these two responses are often treated as related but distinct components of litter processing (Pascoal & Cássio, 2004). Once litter-associated microbes can also use dissolved C from the surrounding water, the location of respiration no longer identifies the C source being respired. This distinction helps explain why respiration and litter decomposition were only partly coupled, and why microbial activity could be associated with DOC concentration without producing a simple, proportional decomposition response.

These separate particulate and dissolved C pathways are especially important to consider in boreal headwaters, where decomposing litter and terrestrially derived DOC enter streams through partly linked but not identical pathways. Riparian vegetation determines the quality and timing of particulate litter inputs, while soil water flow paths, mire cover, hydrological connectivity, and seasonality regulate the concentration and character of DOC reaching the stream channel (Ågren et al., 2008; Laudon et al., 2011). In this experiment, all channels received the same ambient boreal stream water, so the associations of time-adjusted DOC concentration with respiration and relative mass remaining reflected variation in concentration within a shared DOC pool. A possible explanation is that DOC provided an additional resource for litter-associated microorganisms, helping sustain microbial biomass or activity on the litter surface. At a given incubation duration, this could support both greater respiration and greater litter decomposition, even though mass loss slowed over the experiment as readily available litter compounds declined. In previous experiments, additional labile C has been shown to stimulate (prime) leaf litter decomposition under some conditions, although this response is not universal and may depend on nutrient availability, litter type, and decomposer activity (Danger et al., 2013; Soares et al., 2017). This could explain why higher time-adjusted DOC concentration was associated with both higher respiration and greater mass loss, while respiration and decomposition did not change proportionally over time. It is also consistent with isotope-based studies showing that litter-associated microorganisms can assimilate dissolved C from the water column during decomposition (Pastor et al., 2014; Bastias et al., 2020). We cannot exclude, however, that in natural boreal streams, differences in DOC concentration are accompanied by differences in DOC composition and lability, because land cover and landscape configuration strongly influence stream DOC composition (Kothawala et al., 2015; Bastias & Jonsson, 2024), meaning that changes in boreal DOC regimes may affect litter-associated microbial activity through both DOC quantity and DOC quality (Bastias & Jonsson, 2024).

The broader implication of our results is that the relative contribution of litter-derived C and DOC to litter-associated microbial respiration may change as decomposition progresses, while changes in boreal DOC regimes could modify this balance. Under ongoing browning and changing hydrological connectivity, increases in DOC concentration or changes in DOC quality could shift the balance between respiration of litter-derived C and respiration of water column DOC (Kritzberg et al., 2020; Bastias & Jonsson, 2024). Similarly, forestry or buffer strip management could alter the relative inputs of alder, birch, and spruce litter (Hasselquist et al., 2021), while climate-driven shifts in boreal riparian vegetation could further alter the quality of litter entering streams (Nilsson et al., 2013). Alder and birch litter are generally processed faster than spruce litter, although our results show that the two deciduous species are not functionally equivalent. Such changes in litter composition could therefore alter the rate and timing of microbial C processing and the availability of resources to detritivorous macroinvertebrates (Lidman et al., 2017b). Predicting C processing in boreal streams therefore requires distinguishing between decomposition of particulate litter and microbial activity occurring on litter surfaces. A key next step is to identify which microbial groups use DOC during litter decomposition, and for which litter types and during which decomposition stages this becomes important.

## Conclusion

Overall, our results show that litter-associated microbial activity in boreal headwaters cannot be understood only as decomposition of the litter substrate on which the microbes occur. Decomposing litter functions as a biologically active interface where particulate litter C and DOC are processed by microbial communities that change with litter type and decomposition stage. This means that changes in riparian vegetation and DOC regimes may alter not only the rate of in-stream organic matter processing, but also the balance between particulate and dissolved C pathways. In boreal streams, where both litter inputs and DOC export are sensitive to climate, hydrology, and vegetation change, understanding microbial C processing will therefore require distinguishing between where microbial activity occurs, which microbes are involved, and which C source supports that activity. Without resolving this interaction between litter decomposition and DOC use, we cannot fully understand how climate-driven changes in terrestrial C inputs will affect boreal stream C cycling.

## Acknowledgements

We thank Julia Siegel for assistance with fieldwork and laboratory analyses and Bumin Kaan Kiraz for assistance with laboratory analyses. We also thank Meredith Blackburn for leading the design and construction of the experimental mesocosm system used in this study.

## Statements and Declarations

### Authors’ contributions

Matouš Jimel, Lenka Kuglerová, Judith Sarneel, and Micael Jonsson contributed to conceptualization and study design. Matouš Jimel carried out fieldwork, laboratory work, data analysis, visualization, and writing of the original draft. Micael Jonsson, Lenka Kuglerová, and Judith Sarneel contributed to fieldwork and interpretation of the results. Lenka Kuglerová facilitated access to the experimental channels. Eric Capo provided laboratory resources and performed bioinformatic processing of sequence data to generate the ASV tables. All authors reviewed and edited the manuscript and approved the final version.

### Data availability

The data supporting the findings of this study will be deposited in a public repository before publication. Repository links and accession numbers will be added to the final manuscript.

