## Supplementary Material for "Leaf litter decomposition, microbial respiration and community succession under variable dissolved organic carbon availability in boreal streams"

### Supplementary tables

**Table S1 DOC concentration, channel discharge, and calculated DOC flux for each experimental channel at each litter-retrieval date**

DOC concentration was measured at the distribution box level, with each box supplying three channels. DOC concentrations for retrieval days 9 and 50 were estimated by linear interpolation between sampling dates. DOC flux was calculated as DOC concentration × channel discharge.

| Retrieval date | Retrieval day | Distribution box | Channel | DOC concentration (mg L-1) | DOC data | Discharge (L s-1) | DOC flux (mg s-1) |
|---|---|---|---|---|---|---|---|
| 28/08/24 | 2 | 1 | 1 | 47.9 | Measured | 0.40 | 19.27 |
| 28/08/24 | 2 | 1 | 2 | 47.9 | Measured | 0.57 | 27.26 |
| 28/08/24 | 2 | 1 | 3 | 47.9 | Measured | 0.37 | 17.84 |
| 28/08/24 | 2 | 2 | 4 | 47.9 | Measured | 0.36 | 17.42 |
| 28/08/24 | 2 | 2 | 5 | 47.9 | Measured | 0.37 | 17.55 |
| 28/08/24 | 2 | 2 | 6 | 47.9 | Measured | 0.12 | 5.72 |
| 28/08/24 | 2 | 3 | 7 | 48.1 | Measured | 0.13 | 6.35 |
| 28/08/24 | 2 | 3 | 8 | 48.1 | Measured | 0.09 | 4.43 |
| 28/08/24 | 2 | 3 | 9 | 48.1 | Measured | 0.11 | 5.21 |
| 28/08/24 | 2 | 4 | 10 | 48.6 | Measured | 0.08 | 3.79 |
| 28/08/24 | 2 | 4 | 11 | 48.6 | Measured | 0.64 | 31.07 |
| 28/08/24 | 2 | 4 | 12 | 48.6 | Measured | 0.62 | 30.38 |
| 4/9/2024 | 9 | 1 | 1 | 44.99 | Interpolated | 0.59 | 26.58 |
| 4/9/2024 | 9 | 1 | 2 | 44.99 | Interpolated | 0.71 | 32.14 |
| 4/9/2024 | 9 | 1 | 3 | 44.99 | Interpolated | 0.56 | 25.20 |
| 4/9/2024 | 9 | 2 | 4 | 45.03 | Interpolated | 0.06 | 2.57 |
| 4/9/2024 | 9 | 2 | 5 | 45.03 | Interpolated | 0.41 | 18.27 |
| 4/9/2024 | 9 | 2 | 6 | 45.03 | Interpolated | 0.06 | 2.84 |
| 4/9/2024 | 9 | 3 | 7 | 44.95 | Interpolated | 0.06 | 2.82 |
| 4/9/2024 | 9 | 3 | 8 | 44.95 | Interpolated | 0.44 | 19.60 |
| 4/9/2024 | 9 | 3 | 9 | 44.95 | Interpolated | 0.66 | 29.54 |
| 4/9/2024 | 9 | 4 | 10 | 45.8 | Interpolated | 0.53 | 24.29 |
| 4/9/2024 | 9 | 4 | 11 | 45.8 | Interpolated | 0.53 | 24.11 |
| 4/9/2024 | 9 | 4 | 12 | 45.8 | Interpolated | 0.43 | 19.91 |
| 17/09/24 | 22 | 1 | 1 | 39.6 | Measured | 0.65 | 25.55 |
| 17/09/24 | 22 | 1 | 2 | 39.6 | Measured | 0.46 | 18.30 |
| 17/09/24 | 22 | 1 | 3 | 39.6 | Measured | 0.65 | 25.79 |

| | | | | | | | |
|---|---|---|---|---|---|---|---|
| 17/09/24 | 22 | 2 | 4 | 39.7 | Measured | 0.43 | 17.21 |
| 17/09/24 | 22 | 2 | 5 | 39.7 | Measured | 0.53 | 20.97 |
| 17/09/24 | 22 | 2 | 6 | 39.7 | Measured | 0.46 | 18.28 |
| 17/09/24 | 22 | 3 | 7 | 39.1 | Measured | 0.84 | 32.98 |
| 17/09/24 | 22 | 3 | 8 | 39.1 | Measured | 0.62 | 24.33 |
| 17/09/24 | 22 | 3 | 9 | 39.1 | Measured | 0.77 | 29.91 |
| 17/09/24 | 22 | 4 | 10 | 40.6 | Measured | 0.68 | 27.73 |
| 17/09/24 | 22 | 4 | 11 | 40.6 | Measured | 0.75 | 30.56 |
| 17/09/24 | 22 | 4 | 12 | 40.6 | Measured | 0.11 | 4.38 |
| 15/10/24 | 50 | 1 | 1 | 35.3 | Interpolated | 0.11 | 4.03 |
| 15/10/24 | 50 | 1 | 2 | 35.3 | Interpolated | 0.17 | 5.98 |
| 15/10/24 | 50 | 1 | 3 | 35.3 | Interpolated | 0.04 | 1.42 |
| 15/10/24 | 50 | 2 | 4 | 35.53 | Interpolated | 0.10 | 3.39 |
| 15/10/24 | 50 | 2 | 5 | 35.53 | Interpolated | 0.71 | 25.25 |
| 15/10/24 | 50 | 2 | 6 | 35.53 | Interpolated | 0.41 | 14.72 |
| 15/10/24 | 50 | 3 | 7 | 33.89 | Interpolated | 0.62 | 20.99 |
| 15/10/24 | 50 | 3 | 8 | 33.89 | Interpolated | 0.54 | 18.46 |
| 15/10/24 | 50 | 3 | 9 | 33.89 | Interpolated | 0.52 | 17.57 |
| 15/10/24 | 50 | 4 | 10 | 35.59 | Interpolated | 0.08 | 2.89 |
| 15/10/24 | 50 | 4 | 11 | 35.59 | Interpolated | 0.10 | 3.42 |
| 15/10/24 | 50 | 4 | 12 | 35.59 | Interpolated | 0.08 | 2.95 |
| 30/10/24 | 65 | 1 | 1 | 33 | Measured | 0.08 | 2.66 |
| 30/10/24 | 65 | 1 | 2 | 33 | Measured | 0.20 | 6.58 |
| 30/10/24 | 65 | 1 | 3 | 33 | Measured | 0.57 | 18.93 |
| 30/10/24 | 65 | 2 | 4 | 33.3 | Measured | 0.20 | 6.81 |
| 30/10/24 | 65 | 2 | 5 | 33.3 | Measured | 0.51 | 16.83 |
| 30/10/24 | 65 | 2 | 6 | 33.3 | Measured | 0.68 | 22.63 |
| 30/10/24 | 65 | 3 | 7 | 31.1 | Measured | 0.61 | 18.85 |
| 30/10/24 | 65 | 3 | 8 | 31.1 | Measured | 0.57 | 17.70 |
| 30/10/24 | 65 | 3 | 9 | 31.1 | Measured | 0.61 | 18.85 |
| 30/10/24 | 65 | 4 | 10 | 32.9 | Measured | 0.23 | 7.70 |
| 30/10/24 | 65 | 4 | 11 | 32.9 | Measured | 0.58 | 19.19 |
| 30/10/24 | 65 | 4 | 12 | 32.9 | Measured | 0.80 | 26.17 |

**Table S2 Tukey-adjusted pairwise contrasts among litter types for relative mass remaining at each retrieval date, estimated from the final linear mixed effects model**

Contrasts are shown as first litter type minus second litter type. Estimates were calculated with time-adjusted DOC concentration held at its mean value, corresponding to DOC concentrations with no positive or negative deviation from the seasonal DOC trend for a given incubation time.

| Retrieval day | Contrast | Estimate | SE | df | t-ratio | Tukey-adjusted p |
|---|---|---|---|---|---|---|
| 2 | Alder − Birch | −0.0158 | 0.0163 | 156 | −0.970 | 0.597 |
| 2 | Alder − Spruce | −0.0430 | 0.0165 | 156 | −2.607 | 0.027 |
| 2 | Birch − Spruce | −0.0271 | 0.0165 | 156 | −1.645 | 0.230 |
| 9 | Alder − Birch | −0.0205 | 0.0141 | 156 | −1.455 | 0.315 |
| 9 | Alder − Spruce | −0.0366 | 0.0143 | 156 | −2.564 | 0.030 |
| 9 | Birch − Spruce | −0.0161 | 0.0143 | 156 | −1.125 | 0.500 |
| 22 | Alder − Birch | −0.0293 | 0.0113 | 156 | −2.589 | 0.028 |
| 22 | Alder − Spruce | −0.0248 | 0.0114 | 156 | −2.173 | 0.079 |
| 22 | Birch − Spruce | 0.0045 | 0.0114 | 156 | 0.395 | 0.918 |
| 50 | Alder − Birch | −0.0481 | 0.0144 | 156 | −3.331 | 0.003 |
| 50 | Alder − Spruce | 0.0007 | 0.0142 | 156 | 0.050 | 0.999 |

| Retrieval day | Contrast | Estimate | SE | df | t-ratio | Tukey-adjusted p |
|---|---|---|---|---|---|---|
| 50 | Birch − Spruce | 0.0488 | 0.0144 | 156 | 3.398 | 0.003 |
| 65 | Alder − Birch | −0.0582 | 0.0196 | 156 | −2.972 | 0.010 |
| 65 | Alder − Spruce | 0.0144 | 0.0192 | 156 | 0.748 | 0.735 |
| 65 | Birch − Spruce | 0.0726 | 0.0195 | 157 | 3.727 | 0.001 |

### Supplementary figures

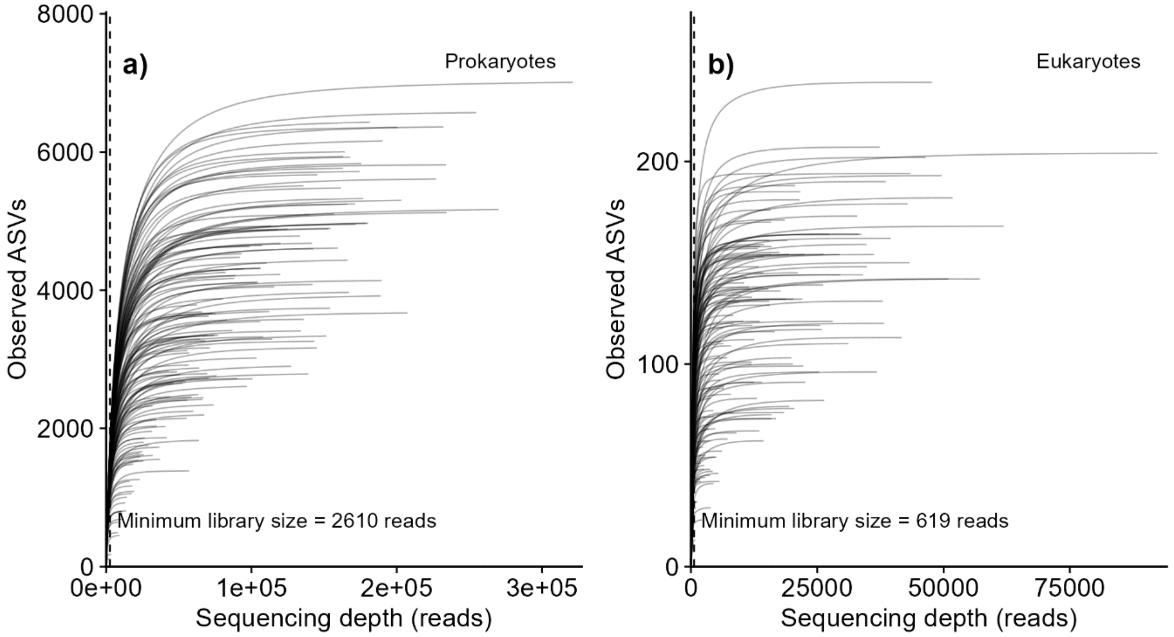


**Fig. S1** Rarefaction curves for final prokaryotic (a) and eukaryotic (b) ASV inventories after ASV table cleaning. Curves show observed ASV richness per retained biological sample as a function of sequencing depth. Dashed vertical lines indicate the minimum retained library size in each inventory, 2,610 reads for the prokaryotic inventory and 619 reads for the eukaryotic inventory

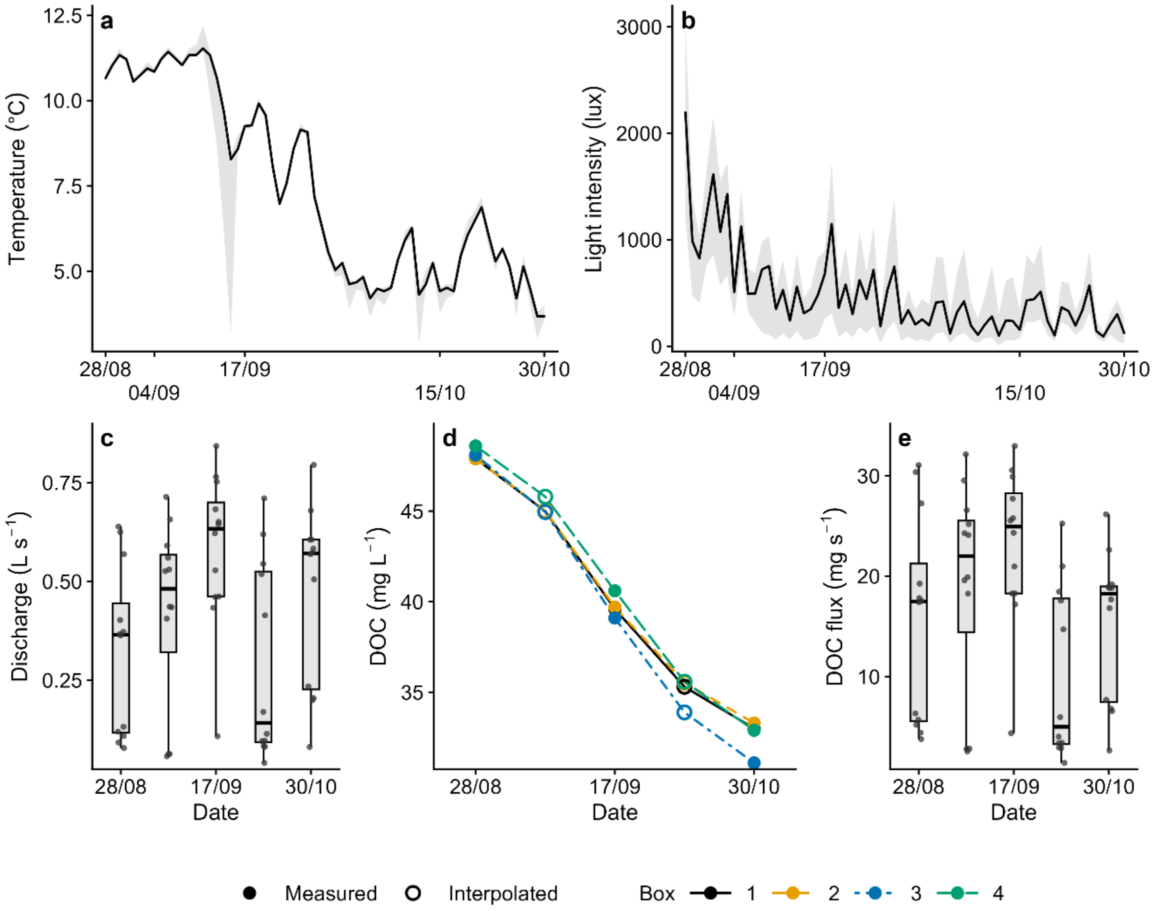


**Fig. S2** Environmental conditions during the experiment. (a) Daily median stream temperature and (b) daytime light intensity across channels, with ribbons showing minimum to maximum ranges. (c) Discharge across retrieval dates. (d) DOC concentration in the four distribution boxes, with filled and open symbols showing measured and interpolated values. (e) DOC flux across retrieval dates, calculated as discharge × DOC concentration. Points in (c) and (e) represent individual channels
